# CamelliA-based simultaneous imaging of Ca^2+^ dynamics in subcellular compartments

**DOI:** 10.1101/2021.05.31.446496

**Authors:** Jingzhe Guo, Jiangman He, Katayoon Dehesh, Zhenbiao Yang

**Author notes:** Author contributions: J.G and Z.Y designed the research; J.G and J.H conducted the experiments; J.G wrote the original draft; Z.Y and K.D supervised the research and revised the manuscript; all authors have read and approved the manuscript. Funding information: This work is partly supported by the National Institute of Health (R01GM107311-8) grant awarded to K.D. Corresponding author: Zhenbiao Yang. The author responsible for contact and ensuring the distribution of materials integral to the findings presented in this article in accordance with the Journal policy described in the Instructions for Authors (http://www.plantphysiol.org) is: Zhenbiao Yang.

## Abstract

As a universal second messenger, calcium (Ca^2+^) transmits specific cellular signals via a spatiotemporal signature generated from its extracellular source and internal stores. Our knowledge of the mechanisms underlying generation of a Ca^2+^ signature is hampered by limited tools enabling simultaneous monitoring of the dynamics of Ca^2+^ levels in multiple subcellular compartments. To overcome the limitation and to further improve spatiotemporal resolutions, here we have assembled a molecular toolset (the CamelliA lines) in Arabidopsis that enables simultaneous and high-resolution monitoring of Ca^2+^ dynamics in multiple subcellular compartments through imaging analyses of different single-colored GECIs (Genetically Encoded Calcium Indicators). Indeed, the uncovering of the previously unrecognized Ca^2+^ signatures in three types of Arabidopsis cells in response to internal and external cues is a testimony to the wide applicability of the newly generated toolset for elucidating the subcellular sources contributing to the Ca^2+^signatures in plants.

**One sentence summary:** A toolset for simultaneous imaging of Ca^2+^ dynamics in subcellular compartments has uncovered unrecognized Ca^2+^ signatures in Arabidopsis cells in response to developmental and external cues.

## Introduction

As second messengers, specific Ca^2+^ signatures exhibit as spikes, waves, and oscillations with a defined duration, amplitude, frequency, and/or subcellular location to regulate plant adaptive responses to developmental and environmental signals (Dodd et al., 2010; Kudla et al., 2018; Vigani and Costa, 2019; Lamers et al., 2020). Ca^2+^ signatures are generated by Ca^2+^ channels, Ca^2+^ pumps, and/or Ca^2+^ transporters localized in the plasma membrane or organellar membranes (Stael et al., 2012; Costa et al., 2018; Kudla et al., 2018; Pan et al., 2019; Hilleary et al., 2020; He et al., 2021), and presumably decoded by Ca^2+^ sensing proteins, e.g., calmodulin, calmodulin-like proteins, calcium-dependent protein kinases (CDPKs), and calcineurin B-like proteins (CBLs) (Harmon et al., 2000; Cheng et al., 2002; Luan et al., 2002; Harper et al., 2004; McCormack et al., 2005; DeFalco et al., 2009; Weinl and Kudla, 2009; Kudla et al., 2018; Tang et al., 2020). The mechanisms for encoding Ca^2+^ signatures are poorly characterized, largely due to the spatiotemporal complexity of Ca^2+^ signaling, which often involves a large number of Ca^2+^ signature encoders and decoders distributed widely in multiple subcellular compartments (Stael et al., 2012; Costa et al., 2018; Kudla et al., 2018; Wudick et al., 2018). Untangling the complexity requires tools for simultaneous visualization of Ca^2+^ dynamics in multiple subcellular compartments with high spatiotemporal resolution.

Many Ca^2+^ probes have been developed, including Ca^2+^ selective vibrating probes (Kuhtreiber and Jaffe, 1990; Pierson et al., 1994; Shipley and Feijó, 1999), Ca^2+^ sensitive fluorescent dyes (Grynkiewicz et al., 1985; Minta et al., 1989), and genetically encoded calcium indicators (GECIs) (Shimomura et al., 1962; Miyawaki et al., 1999; Nakai et al., 2001; Zhao et al., 2011). The vibrating probe has a relatively poor spatial resolution, while the fluorescent dyes require extra loading steps (Takahashi et al., 1999; Bothwell et al., 2006). The GECIs provide consistent and convenient means to monitor *in vivo* Ca^2+^ levels with excellent spatiotemporal resolutions, albeit slower kinetics and lower detection efficiency compared to fluorescent dyes (Hendel et al., 2008; Lock et al., 2015). Recent efforts have greatly improved kinetics and sensitivities of GECI-based Ca^2+^ sensors. There are four types of GECIs (Mank and Griesbeck, 2008; Perez Koldenkova and Nagai, 2013) (Supplemental Table S1). The aequorin bioluminescent Ca^2+^ sensors (Johnson et al., 1995; Sedbrook et al., 1996; Logan and Knight, 2003; Mehlmer et al., 2012; Zhu et al., 2013; Sello et al., 2018) are excellent for accurate quantification of Ca^2+^ level at the tissue level, but inadequate for high spatial resolution imaging at subcellular levels (Zhu et al., 2013; Xiong et al., 2014; Sello et al., 2018). The FRET-based ratiometric Ca^2+^ sensors, such as YC3.6 (Krebs et al., 2012; Loro et al., 2012), YC4.6 (Loro et al., 2016), D3cpv (Loro et al., 2013), D4ER (Bonza et al., 2013), *etc.*, offer good spatial resolution at subcellular levels, absolute quantification of Ca^2+^ levels, and tolerance to pH fluctuations, but have a poor dynamic range and require two channels imaging, limiting their applications in simultaneous imaging in multicellular compartments. The non-FRET-based ratiometric Ca^2+^ sensors, such as GCaMP-R (Cho et al., 2017), MatryoshCaMP6s (Ast et al., 2017) family Ca^2+^ sensors, and R-GECO1-mTurquoise sensor (Waadt et al., 2017), retain the advantages of the FRET-based Ca^2+^ sensors, and have dramatically increased dynamic ranges, but are still limited in multiple channel imaging. The single-colored GECIs, such as R-GECO1 (**r**ed fluorescent, **g**enetically-**e**ncoded **C**a^2+^ indicator for **o**ptical imaging), consist of a calmodulin (CaM) domain, a CaM-binding domain from myosin light chain kinase (M13), and a circularly permuted fluorescent protein (cpFP) (Perez Koldenkova and Nagai, 2013). Upon Ca^2+^ binding, CaM interacts with CaM binding domain, resulting in a conformational change in cpFP and increased fluorescence (Mank and Griesbeck, 2008). The GECIs offer a higher dynamic range and kinetic superiority to the ratiometric GECIs. Importantly, the diverse spectral properties render them ideal for imaging Ca^2+^ dynamics in multiple subcellular compartments (Whitaker, 2010; Keinath et al., 2015). Single-colored GECIs have been successfully applied to imaging Ca^2+^ dynamics in plant cells (Costa and Kudla, 2015; Keinath et al., 2015; Waadt et al., 2017; Walia et al., 2018; De Vriese et al., 2019; Marhavy et al., 2019), but have only been used for simultaneous Ca^2+^ imaging in two subcellular compartments (Kelner et al., 2018; Luo et al., 2020).

Here we assembled a toolset, coined as **CamelliA** for simultaneous **<u>Ca</u>**^2+^ imaging across **<u>m</u>**ultiple subc**<u>ell</u>**ular compartments **i**n **<u>A</u>**rabidopsis, for simultaneous imaging of Ca^2+^ dynamics in multiple subcellular compartments with high spatiotemporal resolution. Consequently, we discovered unprecedented rapid cytosolic Ca^2+^ oscillations in pollen tubes and an unrecognized propagating blast of subcellular Ca^2+^ waves in leaf epidermal cells. Hence by enabling simultaneous Ca^2+^ imaging in multiple compartments, **CamelliA** will uncover new Ca^2+^ signatures and allow dissecting the underlying mechanisms of Ca^2+^-mediated signal integration in plant cells.

## Results

### Design and generation of multi-compartmental Ca^2+^ imaging toolset

The design of multi-compartmental Ca^2+^ imaging is based on the expression of targeted GECIs with distinct emissions in various subcellular compartments, including the cytosol, the nucleus, the sub-PM region, the ER lumen, and the chloroplast stroma. Different subcellular compartments display distinct levels of free Ca^2+^ ion (Suzuki et al., 2016; Costa et al., 2018). As such, effective monitoring of the changes in Ca^2+^ levels require the selection of GECIs with appropriate binding affinity to Ca^2+^ (K_d_). Thus, we employed three single-colored GECIs: B-GECO1 (Zhao et al., 2011), GCaMP6f (Chen et al., 2013; Li et al., 2019), and jRGECO1a (Dana et al., 2016), for imaging the cytosolic and the nuclear Ca^2+^ levels, because their K_d_ is within the range of the cytosolic and nuclear Ca^2+^ concentrations (van Der Luit et al., 1999; Mithofer and Mazars, 2002; Stael et al., 2012; Sello et al., 2016; Costa et al., 2018; Jiang et al., 2019). They were each tagged with an N-terminal nuclear export signal peptide from PKIα (Wen et al., 1995) or C-terminal three tandem repeats of the nuclear localization signal from the SV40 large T antigen (Kalderon et al., 1984) for cytosolic and nuclear targeting respectively. To monitor extracellular Ca^2+^ influx through the PM, we used a similar approach as previously described (Krebs et al., 2012; Iwano et al., 2015) by anchoring GCaMP6f and jRGECO1a to the cytosolic side of the PM through C-terminal fusion with AtLTI6b (Cutler et al., 2000). For visualizing Ca^2+^ dynamics in the ER lumen, we selected two low-affinity single-colored GECIs, G-CEPIA1er and R-CEPIA1er (Suzuki et al., 2014) with optimal K_d_ for analyzing Ca^2+^ levels in the ER lumen (Iwano et al., 2009; Stael et al., 2012; Bonza et al., 2013; Costa et al., 2018). ER targeting signal peptide from Arabidopsis AtCRT1A (Bonza et al., 2013) and a KDEL ER-retention signal peptide (Denecke et al., 1992) were fused to N- and C-termini of the GECIs respectively to target them to ER lumen. Lastly, we chose GCaMP6f for sensing the Ca^2+^ level in the chloroplast stroma because YC3.6, a Ca^2+^ sensor with similar Kd, has been successfully applied to assessing Ca^2+^ dynamics in the stroma (Loro et al., 2016). Thus, the N-terminus of GCaMP6f was fused with tandem repeats of chloroplast stroma targeting peptide from Arabidopsis β-amylase 4 (Fulton et al., 2008; Loro et al., 2016) for targeting to the chloroplast stroma. The Kd of the chosen GECIs and the Ca2+ level in the above compartments were summarized in Supplemental Table S2.

To allow Ca^2+^ imaging in various tissues and cell types, all of the above GECIs were driven by the Arabidopsis *UBQ10* promoter, a ubiquitous promoter active in various cell types and tissues including pollen tubes, hypocotyls, roots, cotyledons, true leaves (Geldner et al., 2009; Grefen et al., 2010; Krebs et al., 2012). The nomenclatures and the respective detailed construct designs are summarized in Figure 1 and Supplemental Table S3.

**Figure 1.**
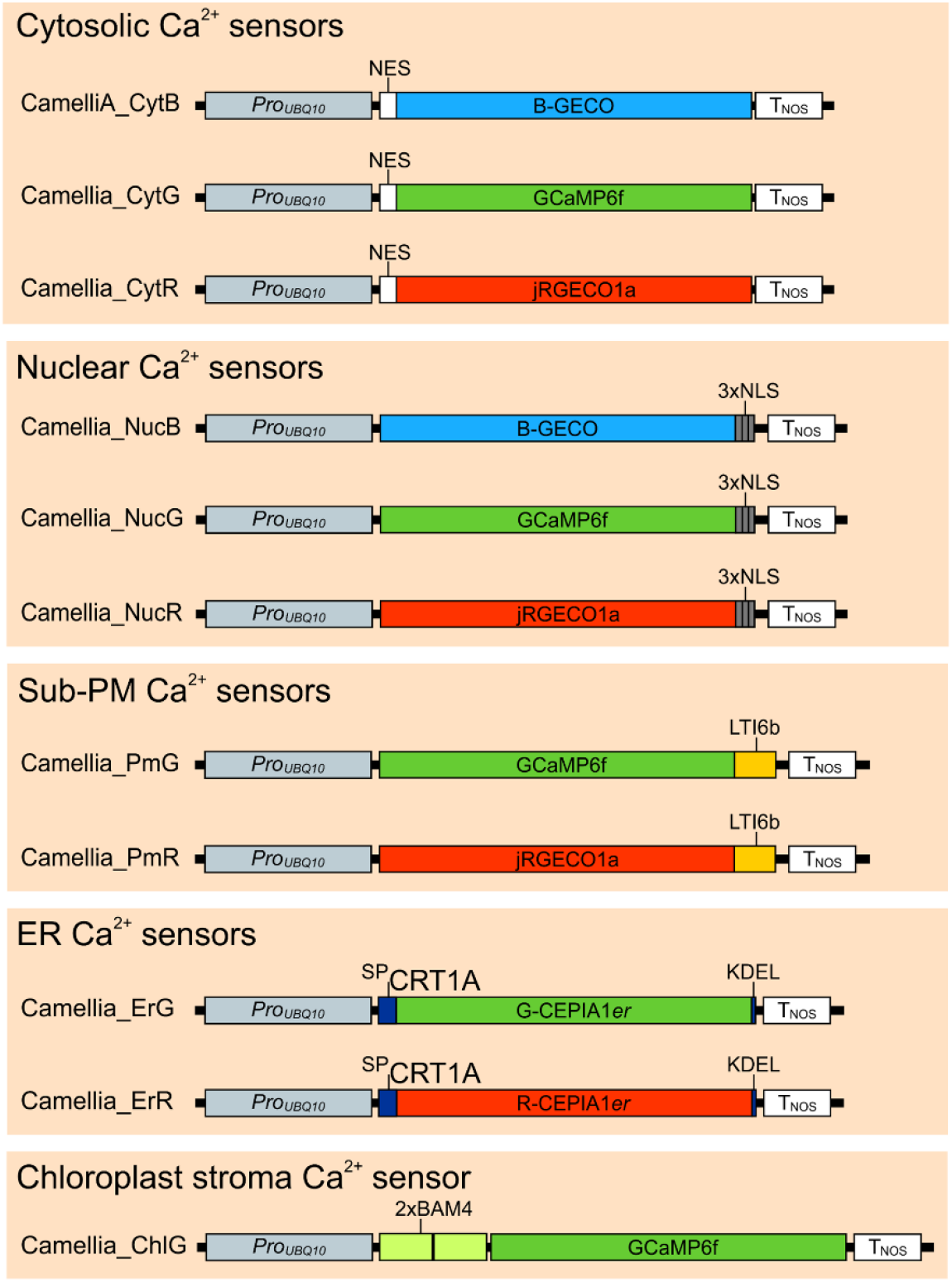
Schematics of various organelle-specific CamelliA Ca^2+^ sensor constructs. Arabidopsis transgenic lines expressing organelle-specific CamelliA calcium sensor constructs under the control of UBIQUITIN 10 (UBQ10) promoter were generated. NES, nuclear export signal; NLS, nuclear localization signal; LTI6b, Arabidopsis LOW TEMPERATURE INDUCED PROTEIN 6B; SP_CRT1A_, the signal peptide of Arabidopsis CALRETICULIN1A; 2xBAM4, two tandem repeats of the signal peptide of Arabidopsis BETA-AMYLASE 4 (BAM4); T_NOS_, NOS terminator.

To acquire transgenic plants without abnormal growth phenotypes that may result from the expression of single-colored GECIs (Waadt et al., 2017), we subjected the transgenic lines to several rounds of stringent selections by analyzing their genetic and physiological performances (Supplemental Figure S1). All final chosen transgenic lines must have a single T-DNA insertion, low or moderate expression levels of the GECIs to avoid adversely affect by overexpression of the GECIs, and similar growth phenotypes as those of wild-type plants. Details of the analyses are described in Supplemental Method S3.

Confocal imaging confirmed the correct subcellular localization of all GECIs in the CamelliA lines. The cytosolic GECIs were excluded from the nuclei (Figure 2A-D; Supplemental Figure S8A-F), while the nuclear GECIs were exclusively localized within the nuclei (Figure 2E-H; Supplemental Figure S8G-I). The PM-anchored Ca^2+^ sensors (CamelliA_PmG/R) exhibited colocalization with the FM4-64-stained PM (Figure 2I-L; Supplemental Figure S8M-O). The two ER-targeted GECIs (CamelliA_ErG/R) were colocalized with the ER marker AtWAK2-CFP-HDEL (Nelson et al., 2007) (Figure 2M-P; Supplemental Figure S8P-R). The colocalization of the GECI with autofluorescent chlorophylls in the CamelliA_ChlG line confirms the correct targeting to chloroplast stroma (Figure 2Q-T).

**Figure 2.**
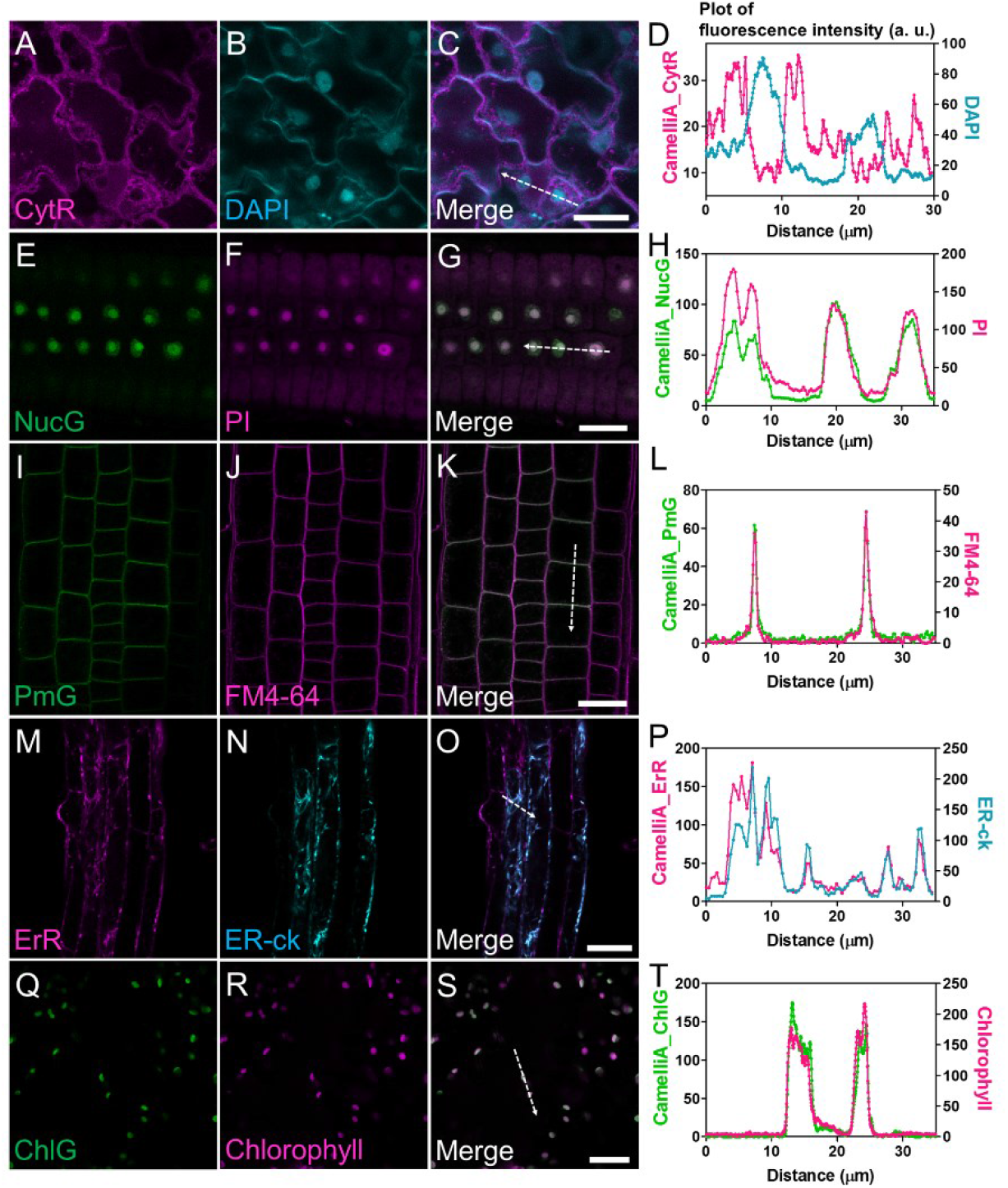
Colocalization of the GECIs and the corresponding subcellular compartments markers in the CamelliA lines. A-D, Representative confocal images of CamelliA_CytR (A), DAPI-stained nuclei (B), and merged image (C). Fluorescence intensity along the direction of the dotted white arrow in the merged image C is plotted in D. E-H, Representative confocal images of CamelliA_NucG (E), PI-stained nuclei (F), and merged image (G). Fluorescence intensity along the direction of the dotted white arrow in the merged image (G) is plotted in H. I-L, Representative confocal images of CamelliA_PmG (I), FM4-64-stained PM (J), and merged image (K). Fluorescence intensity along the direction of the dotted white arrow in the merged image (K) is plotted in L. M-P, Representative confocal images of CamelliA_ErR (M), ER-ck labeled ER (N), and merged image (O). Fluorescence intensity along the direction of the dotted white arrow in the merged image (o) is plotted in P. Q-T, Representative confocal images of CamelliA_ChlG (Q), autofluorescence of the chlorophyll (R), and merged image (S). Fluorescence intensity along the direction of the dotted white arrow in the merged image (S) is plotted in T. Scale bar is 20 µm in C, G, K, S, and 40 µm in O.

### The CamelliA lines robustly report Ca^2+^ signatures in response to extracellular stimuli

To test the performance of the CamelliA lines, we analyzed the spatiotemporal changes in the fluorescence intensity of the GECIs in response to established extracellular stimuli. Extracellular ATP elevates Ca^2+^ levels in many subcellular compartments, including the cytosol (Jeter et al., 2004; Tanaka et al., 2010; Loro et al., 2012), nuclei (Loro et al., 2012; Costa et al., 2013), sub-PM region (Krebs et al., 2012), ER (Bonza et al., 2013), and non-green plastids (Loro et al., 2016) in Arabidopsis.

All three cytosolic Ca^2+^ sensors (CamelliA_CytB/G/R) showed a rapid increase in the cytosolic Ca^2+^ levels with several consecutive peaks of Ca^2+^ after 100 µM ATP application (Figure 3A and B; Supplemental Figure S9A, C), identical to those previously reported using the YC3.6 sensor (Tanaka et al., 2010; Loro et al., 2012). Furthermore, a notable shootward propagation of the cytosolic Ca^2+^ wave was observed (Figure 3B), similar to the shootward Ca^2+^ wave induced by salt treatment (Choi et al., 2014; Evans et al., 2016). The fluorescence intensity of all three cytosolic GECIs did not increase after the mock treatment (Figure 3C; Supplemental Figure S9B and D). The three cytosolic sensors displayed similar peaks of cytosolic Ca^2+^, albeit with some minor differences in their shapes (Figure 3A; Supplemental Figure S9A, C). The differences likely resulted from the different biophysical properties, such as affinities for Ca^2+^ binding (K_d_), response kinetics, and dynamic ranges (Supplemental Table S1). B- GECO1 in the CamelliA_CytB line has a low photostability evidenced by significant photobleaching during the imaging (Supplemental Figure S9B). Fluorescence of the CamelliA_CytG/R lines showed no decrease during the whole period of the mock treatment demonstrating excellent photostability (Figure 3C; Supplemental Figure S9D), and as such these two lines are most suitable for long-term imaging of cytosolic Ca^2+^.

**Figure 3.**
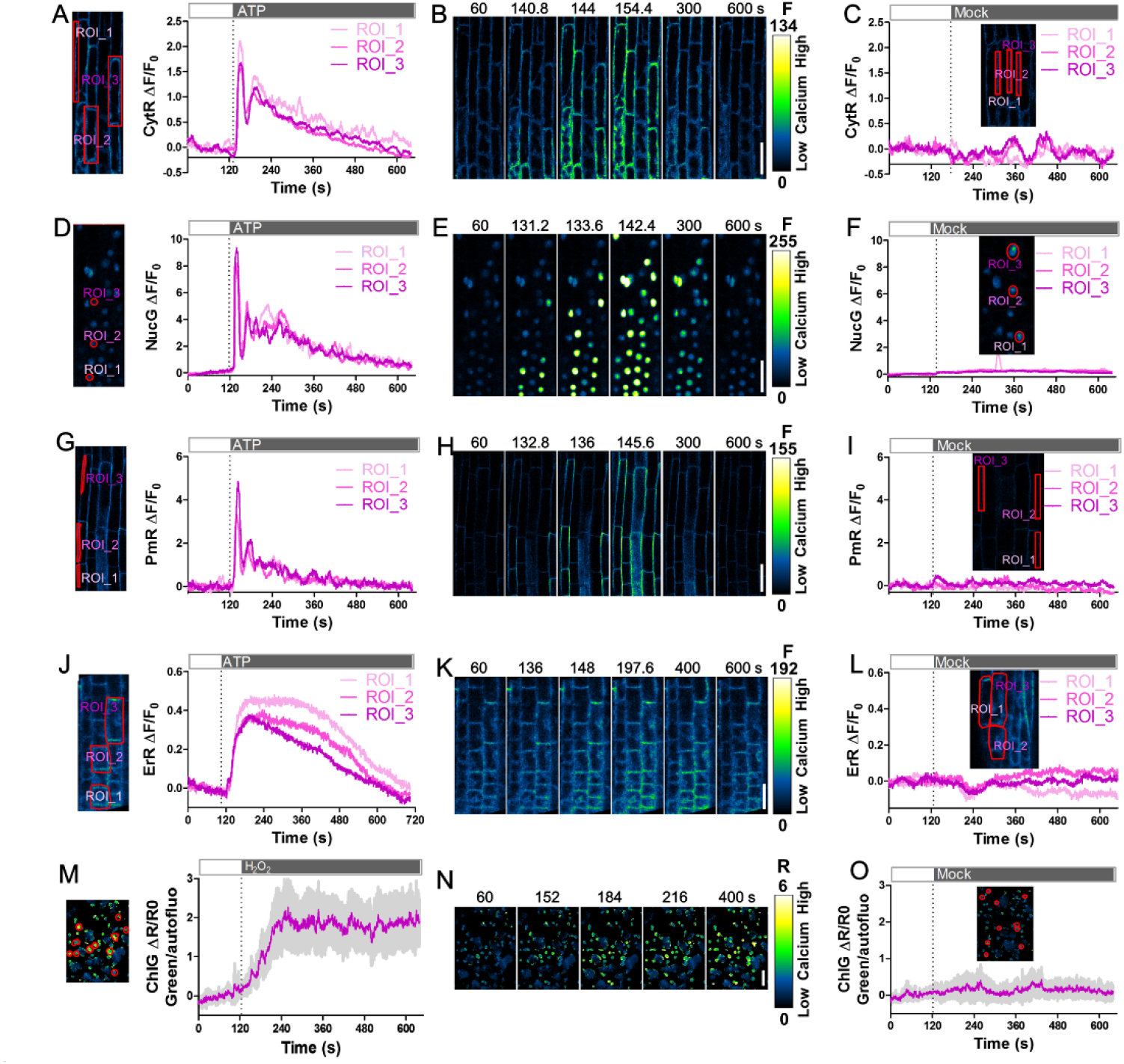
Ca^2+^ signatures at a high spatiotemporal resolution post ATP or H_2_O_2_ treatment of CamelliA lines. A, D, G, J, Time course of fluorescence intensity changes ΔF/F_0_ (right) of the GECIs targeted to four subcellular compartments in the root epidermis of CamelliA_CytR, CamelliA_NucG, CamelliA_PmR, and CamelliA_ErR seedlings before and after extracellular ATP treatment (250 µM ATP for ER-localized GECIs and 100 µM ATP for cytosolic, sub-PM, and nucleus localized GECIs). The labeled ROIs (left) were used for measurement of the GECI intensity in the cytosol (A), the nuclei (D), the sub-PM domain (G), and the ER lumen (J). B, E, H, K, Images of the ATP-treated roots expressing cytosol- (B), nucleus- (E), sub-PM domain- (H), and ER- (K) targeted GECIs at selected time points. C, F, I, L, Time course of fluorescence intensity changes of the GECIs targeted to four subcellular compartments in the root epidermis of CamelliA_CytR, CamelliA_NucG, CamelliA_PmR, and CamelliA_ErR seedlings before and after mock treatment. Inserted images show the ROIs used for measurement of the GECI intensity. M, O, Time course of the change of the ratio ΔR/R_0_ (right) between the stroma-targeted CamelliA_ChlG and the autofluorescence of chlorophyll in cotyledon before and after 10 mM H_2_O_2_ (M) or mock (O) treatment. Mean value (purple line) and 95% confidence intervals (grey shading) were plotted using the data from eleven chloroplasts (left, labeled by red circles). N, Images of the H_2_O_2_- treated cotyledon epidermis at selected time points. Positive fluorescence intensity changes (ΔF/F_0_) and fluorescence intensity ratio changes (ΔR/R_0_) indicate increases in Ca^2+^ concentration in the corresponding subcellular compartment. Representative data from at least three independent biological repeats were shown. Scale bar is 40 µm.

With identical imaging settings, extracellular ATP-induced Ca^2+^ dynamics in the nucleus, the sub-PM region, and the ER were imaged using the CamelliA_NucB/G/R, CamelliA_PmG/R lines, and CamelliA_ErG/R lines, respectively. All nucleus-targeted and PM-anchored GECIs reported several consecutive inductions of Ca^2+^ levels in the nuclei and the sub-PM region, as well as a shootward-propagated Ca^2+^ wave (Figure 3D, G; Supplemental Figure S9E, G, I). No significant changes in the fluorescence intensity were observed in mock treatments (Figure 3F, I; Supplemental Figure S9F, H, J).

Fluorescence of the two ER-targeted GECIs displayed different patterns of changes post ATP treatment (Figure 3J, 4D, and 4E). CamelliA_ErR showed a single transient peak returning to the basal level at the end of the imaging (Figure 3J and 4E), identical to that reported previously using CRT-D4ER sensor (Bonza et al., 2013). In contrast, ErG exhibited multiple peaks (Figure 4D). Moreover, mock treatment triggered a peak in the CamelliA_ErG line, but not in the CamelliA_ErR line (Figure 3L, 4G, and H). Furthermore, G-CEPIA1er used in the ErG line is highly sensitive to pH fluctuations (see details below). These results suggest that CamelliA_ErR serves as an ER Ca^2+^ sensor, whereas the ErG line may not be useful for reporting ER Ca^2+^ signatures.

**Figure 4.**
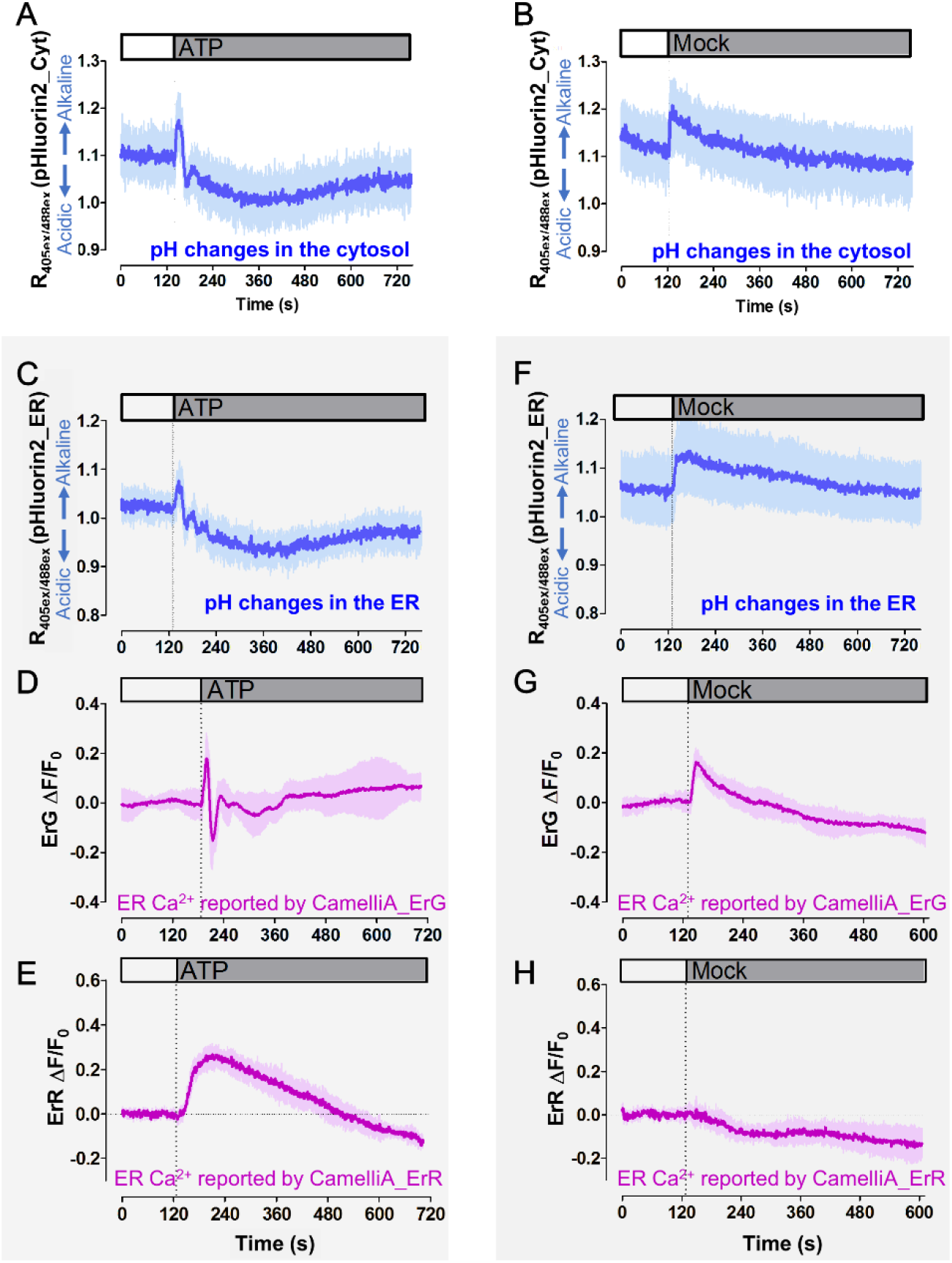
Cytosolic and ER pH fluctuations post extracellular ATP or mock treatment and impact on the ER-targeted GECIs. A-C, F, Monitoring of the pH fluctuations in the cytosol (A, B) and the ER (C, F) of the root epidermis cells treated with 250 µM ATP (A, C) and mock (B, F). Mean (blue line) and 95% confidence interval (blue shading) is calculated from 5 cells from the root epidermis. D-E, G-H, Time course of fluorescence intensity changes (ΔF/F_0_) of the two ER-targeted GECIs, CamelliA_ErG (D, G) and CamelliA_ErR (E, H), in root epidermis with 250 µM ATP treatment (D, E) or mock treatment (G, H). Representative data from at least three independent biological repeats were shown.

We imaged chloroplast stroma GECI in CamelliA_ChlG cotyledons challenged with H_2_O_2_ known to elevate Ca^2+^ in the stroma of chloroplast in leaf cells (Sello et al., 2018). To compensate for the chloroplast movement away from the focal plane resulting in decreased fluorescence, we performed ratio imaging between the fluorescence of the GECI and the autofluorescence of the chlorophyll. Consistent with the previous report (Sello et al., 2018), treatment with 10 mM H_2_O_2_, but not mock treatment, triggered Ca^2+^ elevation in chloroplasts stroma, which reached a plateau in ∼120 seconds post treatment (Figure 3M-O).

In summary, all the CamelliA lines, except CamelliA_ErG, reliably displayed the compartmental specific Ca^2+^ signatures post-treatment with the well-studied extracellular stimuli. Importantly, the CamelliA-based Ca^2+^ imaging showed higher resolution compared with the previous ratiometric Ca^2+^ sensors, because of their enhanced dynamic ranges and no need for ratiometric calculation.

### Most single-colored GECOs in CamelliA lines are insensitive to physiological pH fluctuations under the conditions tested

The concerns that single-colored GECIs are sensitive to pH fluctuations (Keinath et al., 2015) together with the parallel changes in pH and Ca^2+^ levels (Behera et al., 2018), prompted us to examine the influence of pH fluctuations on the GECIs performance in the CamelliA lines. We monitored cellular pH fluctuations using the ratiometric pH sensor, pHluorin2 (Michard et al., 2008; Mahon, 2011). The ratio between the emissions of pHluorin2 excited by 405 nm and 488 nm (R_405ex/488ex_) is proportional to pH values (Mahon, 2011). Free pHluorin2 was used to monitor the pH in both the cytosol and the nucleus, and pHluorin2 fused with AtCRT1a signal peptide and KDEL were used to monitor ER pH. Both constructs are under the Arabidopsis UBQ10 promoter control. Mock treatment with the imaging buffer triggered a transient cytosolic and ER pH elevation (alkalinization) in root epidermal cells (< 10% increase in R_405ex/488ex_) (Figure 4B and F). The buffer application might have triggered mechanosensitive channels to induce ion fluxes through certain membranes and subsequent pH changes (Haswell et al., 2008; Wilson et al., 2013; Peyronnet et al., 2014). However, all the Ca^2+^ sensors, except CamelliA_ErG, showed no changes in fluorescence intensity after mock treatment (Figure3; Supplemental Figure S9). The CamelliA_ErG line exhibited elevated fluorescence intensity after mock treatment (Figure 4G), similar to the pH elevation (Figure 4F). Thus G-CEPIA1er in CamelliA_ErG line is sensitive to pH fluctuations, consistent with the G-CEPIA1er’s highest pKa (8.0) at the Ca^2+^ saturated state compared to all the other GECIs (Supplemental Table S1).

Extracellular ATP triggered similar pH fluctuations both in the cytosol and the ER. Following an initial transient alkalinization, the pH rapidly acidified and then gradually return to the resting level (Figure 4A and C). However, the pH fluctuations did not skew the reported Ca^2+^ signatures in all tested lines, except the CamelliA_ErG line, because they all reported Ca^2+^ signatures consistent with those reported by the less pH-sensitive ratiometric Ca^2+^ sensors. However, G-CEPIA1er again was very sensitive to ATP-induced pH fluctuations (Figure 4D), therefore it is not suitable for reporting Ca^2+^ dynamics in ER.

### CamelliA lines reveal rapid Ca^2+^ oscillations in pollen tubes

Ca^2+^ gradients and oscillations have been extensively characterized in pollen tubes (Pierson et al., 1996; Holdaway-Clarke et al., 1997; Messerli and Robinson, 1997; Michard et al., 2008; Damineli et al., 2017; Barberini et al., 2018). Using ratiometric GECIs such as YC3.6, regular Ca^2+^ oscillations were exclusively observed in non- or slow-growing Arabidopsis pollen tubes, but never in fast-growing tubes (Iwano et al., 2009; Damineli et al., 2017; Diao et al., 2018). The failure to observe Ca^2+^ oscillations in rapidly growing tubes might have resulted from limited temporal resolutions (2-5 seconds) in imaging. Indeed, imaging of CamelliA_CytR pollen tubes at 385 ms per frame revealed Ca^2+^ oscillations at the tip of fast-growing tubes (Figure 5A-C; Supplemental Movie S1). Spectral analyses of the Ca^2+^ oscillations using the intensity vs time showed a major peak at 228 mHz, an indication of Ca^2+^ oscillation with a period of 4.39 s (Figure 5D). Statistical analyses of the cytosolic Ca^2+^ oscillations in 8 pollen tubes, established a mean oscillations period of 5.10 s (Figure 5E). Similar results were also obtained from analyzing the other two cytosolic GECIs in the CamelliA_CytB/G lines (Supplemental Figure S10A-F). Similar Ca^2+^ oscillations were also observed in pollen tubes expressing YC3.6 imaged at 294 ms per frame, albeit with a smaller amplitude and reduced ratio of the FRET channel to CFP channel over time, caused by the photobleaching of YFP (Supplemental Figure S11). These data demonstrated the superiority of the GECIs in the CamelliA lines compared to the ratiometric Ca^2+^ sensors.

**Figure 5.**
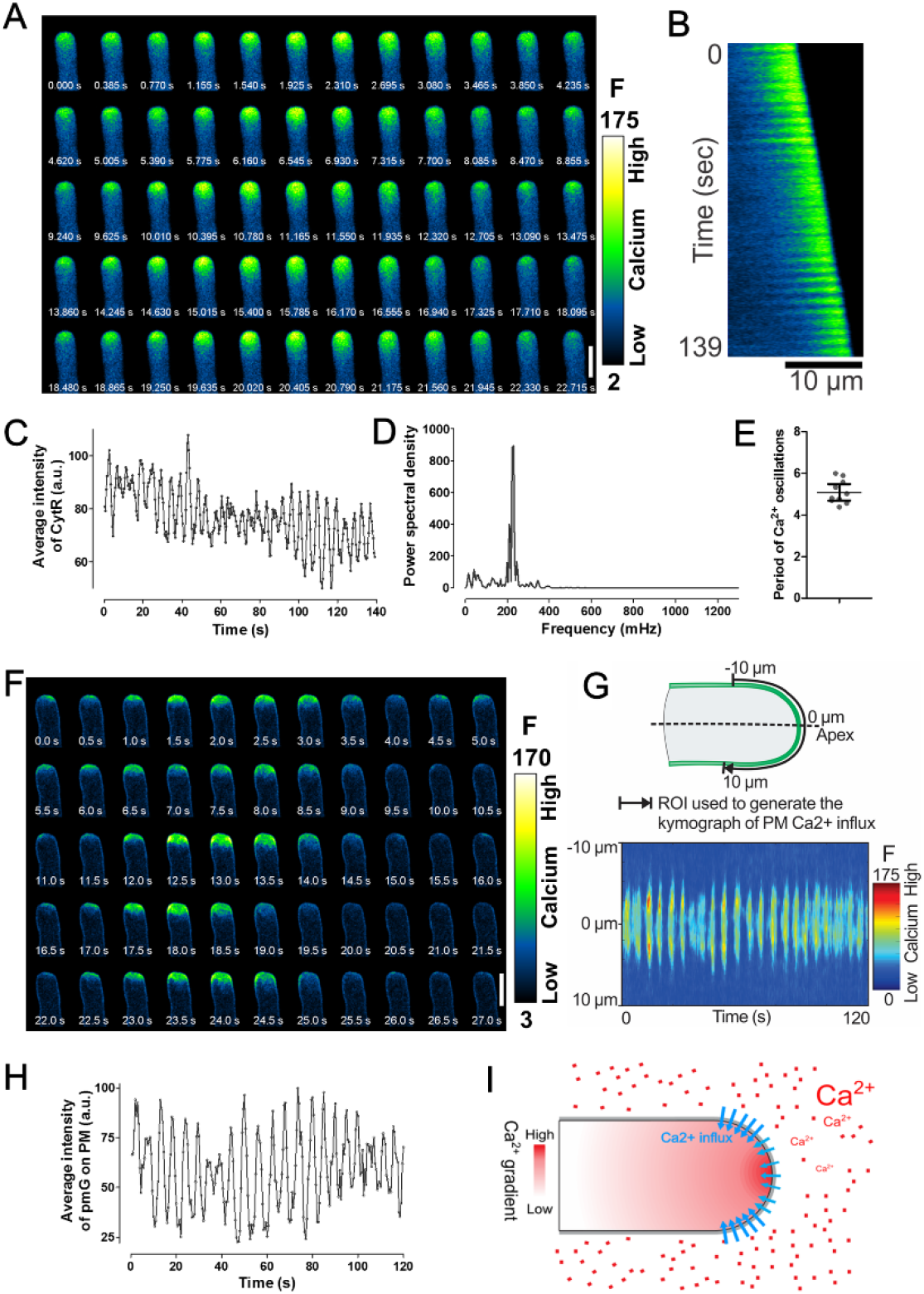
Ca^2+^ imaging at a high spatiotemporal resolution in fast-growing Arabidopsis pollen tubes reveals fast and regular cytosolic Ca^2+^ oscillations and a unique pattern of Ca^2+^ influx on the apical PM. A-B, Time-lapse images (A) and kymograph (B) of a fast- growing pollen tube expressing the cytosolic Ca^2+^ sensor CamelliA_CytR. C-D, Time course of the fluorescence intensity of the CamelliA_CytR in the tip region (6 µm from apex to the shank) of the pollen tube (C) and spectral analysis of the Ca^2+^ oscillations (D). E, Period of the cytosolic Ca^2+^ oscillations from 8 individual fast-growing pollen tubes is plotted in the scatter plot. The mean ± 95% confidence interval is also shown. F, Time-lapse images of a fast-growing pollen tube expressing the PM-anchored Ca^2+^ sensor CamelliA_PmG. G, Kymograph of the CamelliA_PmG fluorescence along the growing PM region of 10 µm on each side of the apex (illustrated in the upper panel) by constantly tracking and analyzing the growing tip using the TipQAD algorithm. H, Time course of the fluorescence intensity of the CamelliA_PmG in the apical PM region (10 µm on each side of the apex) of the pollen tube. I, Model of the Ca^2+^ influx across the apical PM, where the Ca^2+^ influx was initiated at the apex then expanded exclusively to the whole apical PM with stronger activity in the region surrounding the apex. Increases in fluorescence intensity indicate increases in Ca^2+^ concentration. Representative data from at least eight independent biological repeats were shown. Scale bar is 10 µm in A, F.

Proton gradients play an essential role in pollen tube growth (Chen et al., 2020; Hoffmann et al., 2020), and are shown to oscillate (Messerli et al., 1999; Lovy-Wheeler et al., 2006; Damineli et al., 2017; Hoffmann et al., 2020). We assessed whether pH oscillations could contribute to the oscillations of GECI signals in the CamelliA lines using the same pollen tubes co-expressing both CamelliA_CytR and free pHluorin2. The cytosolic pH fluctuated with a small amplitude and showed no overlapping pattern with Ca^2+^ oscillations (Supplemental Figure S12). Furthermore, we imaged Ca^2+^ dynamics in pollen tube co-expressing the cytosolic jRGECO1a and the less pH- sensitive YC3.6. Both sensors displayed an identical pattern of Ca^2+^ oscillations in fast- growing pollen tubes (Supplemental Figure S13) and non-growing pollen tubes (Supplemental Figure S14). These data uncovered rapid oscillations of cytosolic Ca^2+^ with a mean period of 5.10 s in the tip of fast-growing Arabidopsis pollen tubes.

The influx of extracellular Ca^2+^ contributes to the cytosolic Ca^2+^ changes in pollen tubes (Messerli et al., 1999; Hepler et al., 2012; Pan et al., 2019). Ca^2+^ selective vibrating probes detected Ca^2+^ influx across the PM at the extreme apex of lily and poppy pollen tubes (Pierson et al., 1996; Franklin-Tong et al., 2002). However, these probes failed to pinpoint the spatial distribution of the Ca^2+^ influx sites on the PM of pollen tubes. We reasoned that the PM-anchored GECIs may detect the precise sites of Ca^2+^ influxes. Hence, we imaged the pollen tubes of the CamelliA_PmG line at a rate of 2 frames per second. We observed regular oscillations of Ca^2+^ influx across the apical PM of fast-growing pollen tubes (Figure 5F; Supplemental Movie S2). Using TipQAD algorithm for automated tracing of pollen tube tip contour and analyses of fluorescence intensity and distribution on the PM (Tambo et al., 2020), we generated a kymograph along the contour of a 20 µm PM region in the apex of a fast-growing pollen tube (Figure 5G) and a chart of mean intensity vs time over the 20 µm PM region (Figure 5H). Both analyses showed regular oscillations of Ca^2+^ influx across the apical PM with a period of 5.4 s. Furthermore, the kymograph also revealed the influx of Ca^2+^ was exclusively present in the apical PM with a stronger signal in the regions surrounding the extreme apex (Figure 5G). Similar results were also observed using the CamelliA_PmR line (Supplemental Figure S10G-I). Collectively, the data suggest fast and regular oscillations of Ca^2+^ influx at the apical PM of pollen tubes with a period similar to that of the cytosolic Ca^2+^ oscillation.

Since both PM Ca^2+^ influx and cytosolic Ca^2+^ levels oscillate with similar frequencies, we next investigated their temporal relationship by observing these two Ca^2+^ dynamics simultaneously in pollen tubes co-expressing CamelliA_PmG and CamelliA_CytR. As expected, both cytosolic Ca^2+^ and PM Ca^2+^ influx oscillate within the same period (Supplemental Figure S15A-B, D-E). Cross-correlation analyses revealed a slight but notable lag in oscillations of cytosolic Ca^2+^ compared to the PM Ca^2+^ influx (Supplemental Figure S15C, F), supporting the hypothesis that the influx of the Ca^2+^ through the apical PM directly contributes to and drive the elevation of cytosolic Ca^2+^ level.

### High spatiotemporal resolution imaging of multi-compartmental Ca^2+^ dynamics in response to salt treatments

To demonstrate the full capacity of our toolset for multi-compartmental Ca^2+^ imaging, we next imaged Ca^2+^ dynamics in four subcellular compartments, namely the cytosol, the nucleus, the sub-PM region, and the ER, in Arabidopsis root epidermal cells challenged with salt. To avoid any potential adverse impacts on the plant growth as the result of combining all four GECIs, we divided the four sensors into two groups used in independent Ca^2+^ imaging: one group for imaging the Ca^2+^ dynamics in the cytosol, the nucleus, and the sub-PM region (the triple-sensor line generated by crossing CamelliA_CytR, CamelliA_PmG, and CamelliA_NucG lines sequentially); another group for imaging the Ca^2+^ dynamics in the ER and the sub-PM region (the dual-sensor line generated by crossing CamelliA_ErR line to CamelliA_PmG line). The sub-PM Ca^2+^ is shared by both groups, serving as a reference for superimposing the Ca^2+^ dynamics in the four subcellular compartments. The development and growth of homozygous triple- sensor and the dual-sensor lines are indistinguishable from the wild type (Supplemental Figure S5), indicating that co-expressing of the multiple Ca^2+^ sensors have no adverse impacts. To enable fast diffusion of the salt stimulus, the whole seedling was first put on rock wool soaked with 50 µl of imaging buffer, then 150 µl of salt buffer was applied to the seedling (see Supplemental Method S4). Imaging with a sub-second temporal resolution showed that the PM Ca^2+^ influx (sub-PM region) and the cytosolic Ca^2+^ elevated within seconds after salt treatment, followed by the elevation of the nuclear Ca^2+^ a few seconds later (Figure 6A-C), consistent with the previous report (Kelner et al., 2018). Strikingly, a clear propagation of a Ca^2+^ wave on the sub-PM region with a speed of 28.92 ± 12.37 µm/s (mean ± standard deviation) was detected by the PM- anchored Ca^2+^ sensor in the root epidermal cells (Figure 6C; Supplemental Figure S16; Supplemental Movie S3). A similar pattern was also revealed by the cytosolic Ca^2+^ sensor albeit with lower contrast (Figure 6C; Supplemental Movie S3). The elevated nuclear Ca^2+^ was not triggered until the Ca^2+^ wave in the cytosol and sub-PM region reached the nucleus, implying that cytosolic Ca^2+^ elevation may initiate the Ca^2+^ influx into the nucleus (Figure 6C; Supplemental Movie S3).

**Figure 6.**
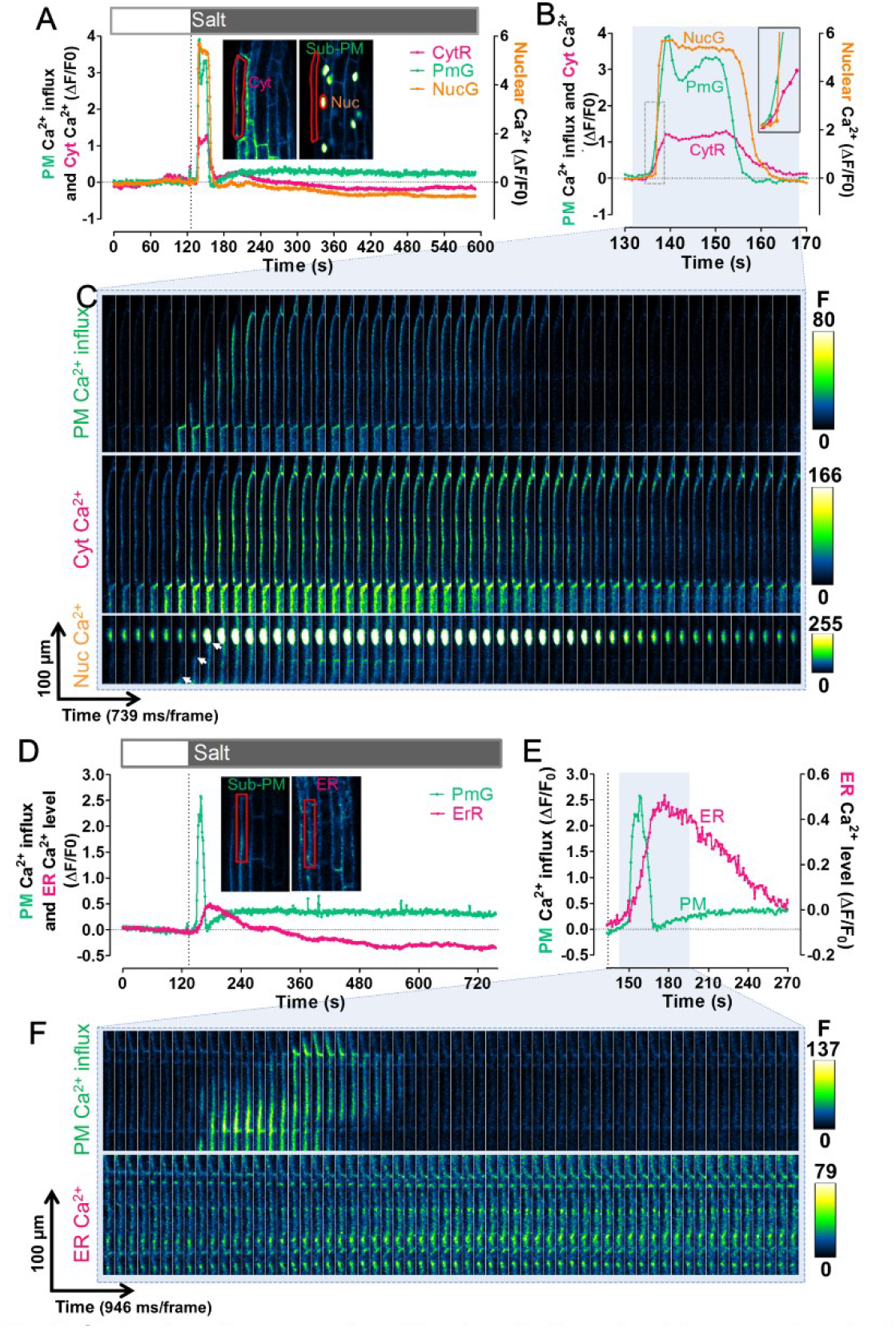
Visualization of the Ca^2+^ signatures in response to salt treatment in four subcellular compartments of the root epidermal cells. A-B, Time course of fluorescent intensity changes (ΔF/F_0_) of the CamelliA_CytR, CamelliA_PmG, and CamelliA_NucG in the same root epidermis cell before and after salt treatment (salt buffer: 150 mM NaCl, 5 mM KCl, 10 mM Ca_2_Cl_2_, 10 mM MES pH5.8). The ROIs highlighted by the red line in the inserted two images were used for quantifying the intensity of corresponding Ca^2+^ sensors. The time course between 130 s and 170 s was replotted in B to show the detail of the Ca^2+^ dynamics in the cytosol, the sub-PM region, and the nucleus. Insert in B showed the detail of the dotted rectangle region. C, Time-lapse images of the sub-PM region (upper), the cytosol (middle), and the nucleus (lower) of the same root epidermis cell expressing the above three Ca^2+^ sensors during the time course of the blue shaded region in B. The white arrows in the lower panel highlight the propagation of the sub-PM Ca^2+^ wave towards the nucleus. D-E, Time course of fluorescent intensity changes (ΔF/F_0_) of the CamelliA_PmG and CamelliA_ErR in the same root epidermis cell before and after salt treatment. The ROIs highlighted by the red line in the inserted two images were used for quantifying the intensity of corresponding Ca^2+^ sensors. The time course between 133 s and 270 s was replotted in E to show the detail of the Ca^2+^ dynamics in the sub-PM region and the ER lumen. F, Time-lapse images of the sub-PM region (upper) and the ER lumen (lower) of the same root cell expressing the two above Ca^2+^ sensors during the time course of the blue shaded region in E. Positive fluorescence intensity changes (ΔF/F_0_) indicate increases of Ca^2+^ concentration in the corresponding subcellular compartment. Representative data from at least three independent biological repeats were shown.

Salt stress also led to the elevation of PM Ca^2+^ influx in the sub-PM region/ER Ca^2+^ dual-sensor line, in a similar pattern observed in the triple-sensor line (Figure 6A-B and D-E; Supplemental Movie S4). However, the elevation of Ca^2+^ level in ER was slower than that in the sub-PM region, reaching the peak when the sub-PM Ca^2+^ restored to the resting level, and then ER Ca^2+^ slowly decreased to below the basal level (Figure 6D-F; Supplemental Movie S4). The mock treatment did not induce detectable changes in the Ca^2+^ levels in the above four compartments (Supplemental Figure S17). Altogether, these observations are in agreement with the previous studies (Knight et al., 1997; Bonza et al., 2013; Corso et al., 2018; Kelner et al., 2018), but differ in that the observations reported here were from the same cell, enabling accurate analysis of the spatiotemporal correlation of the Ca^2+^ dynamics across these four subcellular compartments.

### Subcellular resolution of wounding-induced Ca^2+^ waves

We next expanded our studies to the leaf epidermis, which is commonly used for studying Ca^2+^ responses to biotic and abiotic stresses (Keinath et al., 2015; Vincent et al., 2017; Hilleary et al., 2020). Wounding induces a rapid local elevation of cytosolic Ca^2+^ and initiates a Ca^2+^ wave propagating across tissues (Beneloujaephajri et al., 2013; Nguyen et al., 2018; Toyota et al., 2018), but how the wave is propagated remains unknown. We visualized the subcellular changes in the wounding-triggered Ca^2+^ waves in leaf epidermal cells using our triple-sensor line. The Ca^2+^ levels in all three subcellular compartments of the epidermal cells were monitored after a randomly chosen pavement cell was damaged by laser irradiation. In agreement with previous studies (Beneloujaephajri et al., 2013; Nguyen et al., 2018; Toyota et al., 2018), the cytosolic Ca^2+^ level, as well as the nuclear and sub-PM Ca^2+^ levels, was elevated immediately after the laser-irradiation at the wounding site (Figure 7). Strikingly, a shockwave-like propagation of Ca^2+^ radially spread away from the wounding site in these subcellular compartments (Figure 7A; Supplemental Movie S5). We further analyzed the spatiotemporal pattern of the Ca^2+^ wave in the cytosol and the sub-PM region by generating kymographs on each imaging channel along the edge of the damaged cell to distal cells (Figure 7B). This analysis clearly showed a propagation of Ca^2+^ in both subcellular compartments, from the wounding site to distal cells with an average speed of 9.8 ± 3.1 µm/s (mean ± standard deviation).

**Figure 7.**
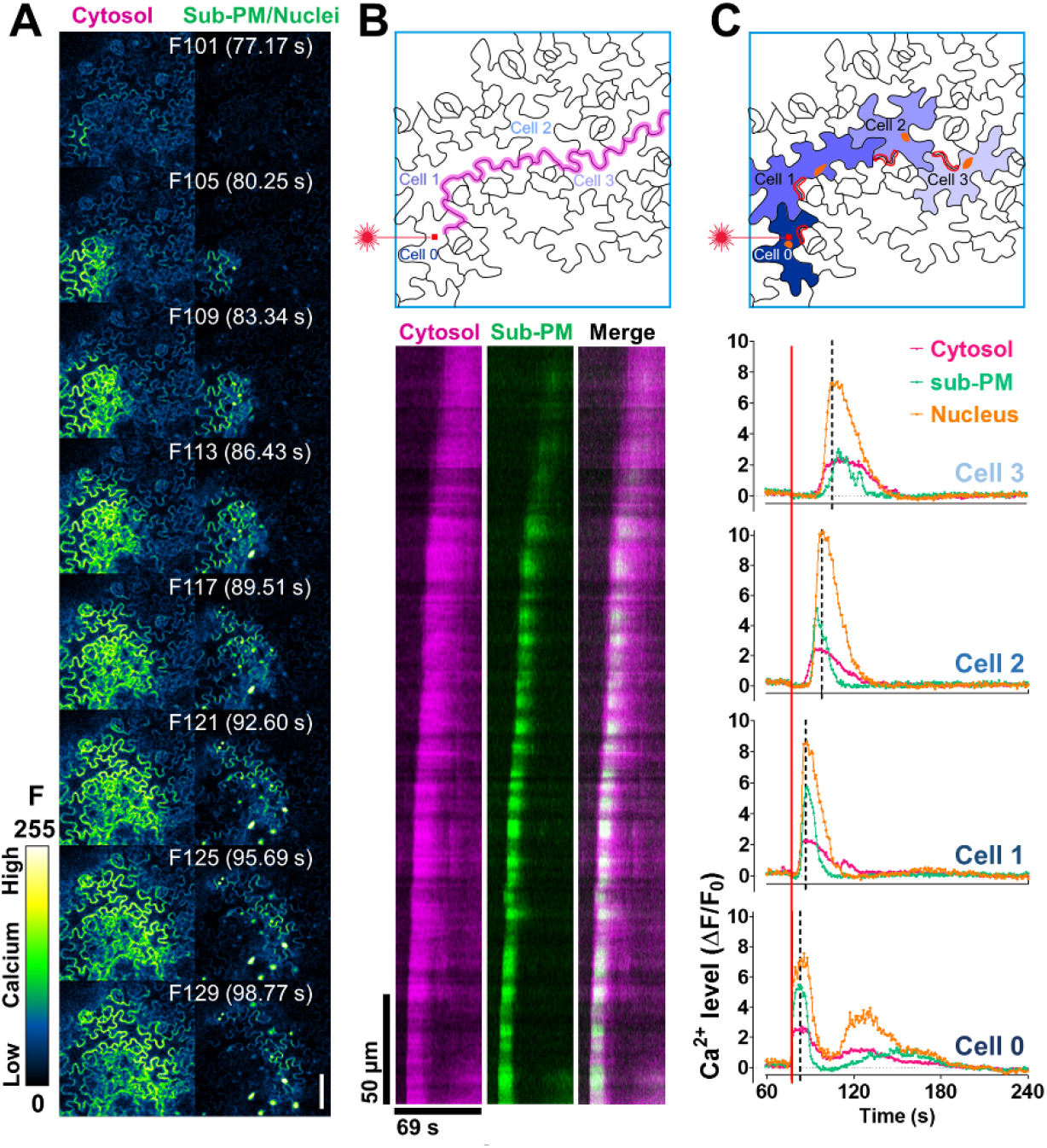
Laser-wounding triggered a local blastwave-like Ca^2+^ wave in the leaf epidermis. A, Time-lapse images of the leaf epidermis after laser-wounding. The cytosolic Ca^2+^ dynamics were shown on the left, and the Ca^2+^ dynamics in the sub-PM regions and the nuclei were shown on the right. The number (F101 - F129) and time of each frame were labeled outside and inside of the parenthesis respectively. At frame 100 (F100), the red square labeled region (upper panel in b and c) of the pavement cell was irradiated by a 405 nm laser. B, Kymograph (lower panel) of the CamelliA_CytR and CamelliA_PmG fluorescence along the path indicated as the purple line across Cell 0-3 in the upper panel. The Ca^2+^ propagated in both cytosol and sub-PM from the wounding site to distal cells with an average speed of 9.8 ± 3.1 µm/s (mean ± standard deviation, achieved by analyzing 8 different Ca2+ propagation directions from the wounding site to its surrounding cells in 4 independent leaves). C, Time course of the fluorescence intensity changes (ΔF/F_0_) of the CamelliA_CytR, CamelliA_PmG, and CamelliA_NucG in four cells (Cell 0 is the wounded cell and Cell 1-3 are three neighboring cells) during the period of 60-240 seconds (F79 - F312). See Supplemental Figure S18 for the time course data of the entire observation. In each cell, the ROI labeled by the red line was used for quantifying the intensity of the Ca^2+^ sensors in the cytosol and sub-PM region, the ROI labeled by the orange spot was used for quantifying the intensity of the nuclear Ca^2+^ sensor. The dotted line in each chart highlights the time point when the nuclear Ca^2+^ level reached its peak. The red square pointed by the laser icon on Cell 0 indicates the wounding site. Positive fluorescence intensity changes (ΔF/F_0_) indicate increases in Ca^2+^ concentration in the corresponding subcellular compartment. Representative data from 10 independent biological repeats were shown. Scale bar is 50 µm in A.

To understand the temporal relationship of the Ca^2+^ dynamics among all three subcellular compartments, we analyzed relative Ca^2+^ levels in these compartments in each of the four chosen cells, which show wave-like Ca^2+^ propagation away from the wounding site (Cell 0) (Figure 7C). In the damaged cell (Cell 0), the Ca^2+^ level in all three compartments increased immediately after the laser irradiation. As a result, we were unable to distinguish their temporal relationships under our current time-lapse imaging (772 ms per frame) (Figure 7C; Supplemental Figure S18). However, in the un- damaged cells, there is an obvious temporal sequence of Ca^2+^ elevation in the compartments. The elevation of cytosolic Ca^2+^ preceded that of sub-PM Ca^2+^ (Ca^2+^ influx) by ∼2-3 seconds, followed by nuclear Ca^2+^ (Figure 7B-C). The elevation of cytosolic Ca^2+^ prior to that the sub-PM Ca^2+^ in un-damaged cells is likely the result of Ca^2+^ release from internal Ca^2+^ storages, such as vacuole or ER (Stael et al., 2012; Costa et al., 2018), though differences in the properties of the reporters in different subcellular localizations could not be excluded. Indeed, several studies supported the contribution of vacuolar Ca^2+^ to cytosolic Ca^2+^ elevations during various biotic and abiotic stresses (Knight et al., 1996; Knight et al., 1997; Kiep et al., 2015; Vincent et al., 2017; Hilleary et al., 2020). In particular, Knight et al (1997) observed a faster Ca^2+^ elevation in the cytosolic microdomain adjacent to the vacuole than that in further away cytosolic regions of osmotically-stressed plants. Thus, mechanical and osmotic stresses apparently trigger a faster efflux of Ca^2+^ from internal stores than the Ca^2+^ influx from the PM in plant cells.

In addition, these subcellular compartments displayed unique Ca^2+^ signatures in all the cells observed. The Ca^2+^ signatures in both the cytosol and the nuclei have longer durations than that of the PM Ca^2+^ elevation, while the nuclear Ca^2+^ signature has higher amplitudes of ΔF/F0 than that of the other two compartments (Figure 7B-C). The distinct Ca^2+^ signatures in the above compartments illustrate the necessity of understanding the Ca^2+^ dynamics in various subcellular compartments when studying the cellular responses to various stresses.

## Discussion

In response to a given environmental and/or developmental signal, a specific spatiotemporal Ca^2+^ signature is produced thereby providing the specificity of calcium signaling. The generation of a unique Ca^2+^ signature typically involves calcium uptakes and releases from multiple subcellular compartments. Therefore, it is essential to visualize Ca^2+^ dynamics in the subcellular compartments simultaneously and within the same cell. In this study, we reported the generation of a toolset, the CamelliA lines, for simultaneously imaging the Ca^2+^ dynamics in multiple subcellular compartments. We first validated the functionality of the Ca^2+^ sensors in our CamelliA lines by using the same well-studied extracellular stimuli and comparing their performance with previously well-established sensors. We also assessed the effects of the pH fluctuations on the performance of our sensor lines. Lastly, we demonstrated the power of our CamelliA lines for visualizing the Ca^2+^ dynamics with high spatiotemporal subcellular resolution in the three most popular cell models in response to environmental stresses and developmental signals, enabling us to uncover several unrecognized Ca^2+^ signatures in plant cells. As such, this toolset will serve as a crucial asset for dissecting Ca^2+^- mediated signaling pathways during stress responses and developmental processes where the Ca^2+^ homeostasis is actively maintained by regulated Ca^2+^ fluxes across multiple subcellular compartments.

### Advantages and consideration of the CamelliA Ca^2+^ sensor toolset

Our toolset offers several key advantages for Ca^2+^ imaging:

**1) Simultaneous imaging of Ca^2+^ dynamics in multiple subcellular compartments**. We demonstrated multiplexed imaging of the Ca^2+^ dynamics in the cytosol, the nuclei, the sub-PM region, and the ER in several types of cells (Figs 5, 6, 7). Although simultaneous imaging of nuclear and cytosolic Ca^2+^ dynamics in plant cells has been reported (Kelner et al., 2018; Luo et al., 2020), our toolset has greatly expanded the choices of GECIs with different colors and targeted various compartments.
**2) Improved spatiotemporal resolution.** Single-colored GECIs have a higher dynamic range than that of ratiometric GECIs (Supplemental Table S1). Less pixel dwell time during confocal imaging is needed to acquire images with a better or equivalent signal-to-noise ratio (SNR) compared with ratiometric GECIs, providing increased temporal resolution. What’s more, the fluorescence decay of GCaMP3 after dissociation with Ca^2+^ is much faster than that of two FRET-based GECIs, TN-XXL, and D3cpV (Tian et al., 2009). GCaMP6f and jRGECO1a that are used in our toolset have even faster kinetics than that of GCaMP3 (Supplemental Table S1). Therefore, GCaMP6f- and jRGECO1a-based sensors in the CamelliA lines are well-suited for resolving temporal details of Ca^2+^ signatures such as fast cytosolic Ca^2+^ oscillations within 5 seconds in Arabidopsis pollen tubes, the most rapid Ca^2+^ oscillations in plants to date. (Figure 5). In addition, the ratio images generated by ratiometric GECIs are noisy, for example, the imaging of the PM Ca^2+^ dynamics with the PM anchored YC3.6 reported noisy Ca^2+^ fluctuations post ATP treatment (Krebs et al., 2012). Our PM-anchored GECIs, instead, have visualized Ca^2+^ influx through PM with minimal noise and a high spatiotemporal resolution (Figure 3G, 5F, 6, 7; Supplemental Figure S9I). Consequently, we uncovered a shockwave-like propagation of Ca^2+^ wave in several subcellular compartments in leaf epidermis after wounding (Figure 7), which was not possible with either YC3.6 (Beneloujaephajri et al., 2013) or aequorin (Kiep et al., 2015).

When using our toolset, one should also consider potential limitations such as determination of absolute Ca^2+^ levels and pH sensitivity and photo-stability for some probes. Absolute quantification of Ca^2+^ concentrations with our toolset is more cumbersome than ratiometric Ca^2+^ sensors and aequorin-based sensors, as the fluorescence intensity of CamelliA sensors depends on the concentration of both Ca^2+^ and Ca^2+^ sensors. Absolute quantification of Ca^2+^ concentration will require calibration for each type of tissue or cell to offset the fluorescence variations resulting from different concentrations of the sensors.

The artificial increase of pH from 6.8 to 8.0 almost doubled the fluorescence intensity of R-GECO1 independent of Ca^2+^ (Keinath et al., 2015). Whereas *in planta* observations showed unnoticeable impacts of the pH fluctuations on the reported Ca^2+^ dynamics (except for the CamelliA_ErG line) (Figure 3 and 4; Supplemental Figure S9, S12). This may be in part due to less drastic fluctuations of pH within the physiological range as compared with manipulated pH from 6.8 to 8.0 (Keinath et al., 2015; Behera et al., 2018). Compared with previous GECIs, the GECIs in our toolset have lower pKa (Supplemental Table S1), which makes them less vulnerable to pH fluctuations. Nevertheless, it is often left for others to check pH fluctuations independently when using our sensors. Finally, B-GECO1 is poorly photostable, which limits the duration of imaging experiments when using B-GECO1 based sensor lines.

### Our CamelliA lines reveal unrecognized characteristics of stimuli-induced Ca^2+^ signatures

Our toolset uncovered several unrecognized features of plant Ca^2+^ signatures. First, we discovered fast oscillations of cytosolic Ca^2+^ and Ca^2+^ influx at the apical PM in fast-growing Arabidopsis pollen tubes. Previous studies failed to reveal these oscillations due to limited temporal resolution (2-5 seconds/frame) (Iwano et al., 2009; Damineli et al., 2017; Diao et al., 2018). Using the PM-anchored Ca^2+^ sensors in the CamelliA_PmG/R lines, we mapped the spatiotemporal pattern of the Ca^2+^ influx across the apical PM of pollen tubes for the first time. Cross-correlation analysis shows Ca^2+^ influxes slightly but notably led (less than 0.5 s) cytosolic Ca^2+^ oscillations, implying that the PM Ca^2+^ influx drives the cytosolic Ca^2+^ oscillations. In contrast, Ca^2+^ specific vibration probes reported presumed Ca^2+^ influxes lagging behind the cytosolic Ca^2+^ oscillations (Holdaway-Clarke et al., 1997; Messerli et al., 1999). The vibration electrode cannot distinguish actual Ca^2+^ influx via the PM from the Ca^2+^ absorbed by the cell wall. Our PM-anchored Ca^2+^ sensors are ideal for visualizing the Ca^2+^ influx from the apoplast to the cytosol.

We also uncovered Ca^2+^ dynamics at sub-second temporal resolution in four subcellular compartments in salt-stressed root epidermal cells and the spatiotemporal relationship of Ca^2+^ fluxes among these compartments. The simultaneous onset of the PM Ca^2+^ influx and cytosolic Ca^2+^ followed by the delayed elevation of the nuclear Ca^2+^ after salt treatment (Figure 6A-C) implies that nuclear Ca^2+^ elevation may be a passive result due to the cytosolic Ca^2+^ increase in response to the salt treatment. Interestingly, the ER Ca^2+^ level starts to increase almost at the same time as the onset of the PM Ca^2+^ influx (Figure 6E and F), implying that uptake of the cytosolic Ca^2+^ by the ER is triggered as soon as the Ca^2+^ influx was triggered.

Last but not the least, we discovered the propagation of a novel Ca^2+^ wave in a shockwave-like pattern initiated from the cell that was damaged by laser irradiation of the leaf epidermis (Figure 7). Although wounding is known to trigger cytosolic Ca^2+^ elevation both locally and systemically (Beneloujaephajri et al., 2013; Kiep et al., 2015; Vincent et al., 2017; Nguyen et al., 2018; Toyota et al., 2018), how wounding-induced Ca^2+^ propagates among cells remains unknown. Our observation showed that laser irradiation on a single pavement cell triggers both intracellular and intercellular shockwave-like Ca^2+^ propagation in all three subcellular compartments, hinting that the intercellular Ca^2+^ propagation is linked to the intracellular Ca^2+^ shockwave. The Ca^2+^ wave propagated in the leaf epidermis with a speed of 9.8 ± 3.1 µm/s, which is similar to the speed of Ca^2+^ wave propagation in nonvascular tissues of aphid-feeding Arabidopsis leaf (5.9 ± 0.6 µm/s) (Vincent et al., 2017) and osmotic-stressed *Physcomitrella patens* plantlet (4.5 ± 3.8 µm/s) (Storti et al., 2018), suggesting a potential common mechanism for Ca^2+^ propagation among non-vascular cells.

We found that the elevation of the cytosolic Ca^2+^ level preceded that of the PM Ca^2+^ influx in non-damaged surrounding cells with a more noticeable time difference in cells further away from the damaged cell (Figure 7B and C). This suggests a contribution from internal Ca^2+^ storage to the cytosolic Ca^2+^ elevation in addition to and faster than PM Ca^2+^ influxes from the apoplast, consistent with the proposal that vacuolar Ca^2+^ storage contributes to the rapid rise of the cytosolic Ca^2+^ during the wounding response (Beyhl et al., 2009; Kiep et al., 2015). The nature of the transmitter(s) of the long-range wounding signal triggering Ca^2+^ release from internal Ca^2+^ storage is unknown. Ca^2+^, glutamate, ROS, and electric signals, which propagate in a wave-like form across tissues, have long been proposed to act as long-range transmitting signals in plants (Miller et al., 2009; Zimmermann et al., 2009; Mousavi et al., 2013; Steinhorst and Kudla, 2013; Choi et al., 2014; Salvador-Recatala et al., 2014; Steinhorst and Kudla, 2014; Nguyen et al., 2018; Toyota et al., 2018; Kumari et al., 2019; Marhavy et al., 2019; Farmer et al., 2020; Zandalinas et al., 2020; Dindas et al., 2021). Our toolset provides a powerful means for testing whether any of these systemic signals play a role in the activation of Ca^2+^ release from internal Ca^2+^ storage.

## Conclusion

In this study, we have assembled a toolset in Arabidopsis, the CamelliA lines, as enabling imaging capability to monitor the dynamics of Ca^2+^ signature simultaneously and with high spatiotemporal resolution in multiple subcellular compartments, including cytosol, sub-PM region, nucleus, ER, and stroma of the chloroplast. Using the toolset, we have identified several previously unrecognized Ca^2+^ signatures in three types of Arabidopsis cells. As such, this toolset will serve as a crucial asset for elucidating the subcellular sources contributing to the Ca^2+^ signature during stress responses and developmental processes. Considering the Arabidopsis UBQ10 promoter can drive ubiquitous expression of YC3.6 in rice (Behera et al., 2015), this toolset shall have a broad application in a number of crop plants.

## Materials and Methods

### Plant materials and growth conditions

All *A. thaliana* lines are in Col-0 grown under a 16-hour-light (22 °C) and 8-hour-dark (19 °C) regime. Seedlings were grown on 1/2 Murashige and Skoog (MS) solid medium (pH=5.7) supplemented with 1% (w/v) sucrose. The seedlings were grown either vertically for 5-6 days to produce roots, or horizontally for 14 days to produce true leaves for Ca^2+^ imaging.

### Chemicals

Stock solutions of chemicals were prepared as followed: ATP (Sigma-Aldrich), FM4-64 (ThermoFisher), and DAPI (Sigma-Aldrich) were dissolved in water to a concentration of 500 mM, 2 mM, and 10 mg/ml respectively. Solution of 1 mg/ml Propidium Iodide (PI) was purchased from ThermoFisher.

### Generation of the CamelliA line**s**

The supplemental information details the generation of the CamelliA constructs (Supplemental Method S1; Supplemental Tables S3, S4), the transformation and screening of the transgenic lines (Supplemental Method S2).

### Ca^2+^ imaging

The wavelengths for the GECI excitation and emission are listed in Supplemental Table S5. Pollen tubes were germinated, mounted, and observed as described (Guo and Yang, 2020). Ca^2+^ imaging in root epidermal cells was performed as described (Behera et al., 2015). A “top-imaging” setup (Keinath et al., 2015) was used to image the Ca^2+^ dynamics in cotyledons and true leaves. Detailed imaging setups are described in Supplemental Method S4.

### Visualization of various subcellular compartments for colocalization analysis

The nuclei were stained with 10 µg/ml PI or DAPI in a solution of 0.05% Tween 20, 1% DMSO, 2 mM CaCl_2_ for 1 hour before observation. PM was stained with 2 µM FM4-64 in MS liquid medium on ice for 5 min and observed immediately. ER marker (ER-ck) (Nelson et al., 2007) was crossed into CamelliA_ErR line for examining their colocalization.

### Image analysis

Images of pollen tubes were analyzed by TipQAD algorithm (Tambo et al., 2020). The temporal relationship between oscillations of the PM Ca^2+^ influx and the cytosolic Ca^2+^ was analyzed using *sample cross-correlation* function in MATLAB. Frequency of the cytosolic Ca^2+^ oscillations was analyzed using *SpectralAnalysis* code (Uhlen, 2004). The fluorescence intensity of root and leaf cells was analyzed in Fiji. Before treatments with ATP, H_2_O_2_, or salt, the resting level of Ca^2+^ in root or leaf samples was imaged for 120 s at sub-second frame rates and plotted as part of the intensity-vs-time plot.

Fluorescence changes (dF/F_0_) of the GECIs in roots and leaves was calculated using the equation of dF/F_0_ = (F_t_ – F_0_)/F_0_, where F_t_ is the fluorescence intensity of the GECIs at a given time point (frame) and F_0_ is the mean fluorescence intensity of the GECIs in all frames when the cells are at resting state. Ratio (between the GCaMP6f intensity and chloroplast autofluorescence) changes (dR/R_0_) of individual chloroplast in H_2_O_2_ treated leaves was calculated using the equation of dR/R_0_ = (R_t_ – R_0_)/R_0_, where Rt is the ratio of the measured chloroplast at a given time point (frame) and R_0_ is the mean ratio of the chloroplast in all frames of the resting state.

### Statistical analysis

The mean and 95% confidence interval of all intensity/ratio vs time data from at least five individual cells were plotted in GraphPad Prism. All data shown are presentative data from at least three biological repeats. The exact number of the cells and biological repeats for analyses is indicated in the legends of corresponding figures.

### Accession numbers

Sequence data from the genes mentioned in this article can be found in the Arabidopsis Genome Initiative under the following accession numbers: CALRETICULIN 1A, AT1G56340; LTI6B, AT3G05890; UBQ10, AT4G05320; BAM4, AT5G55700

## Acknowledgments

We thank members of the Yang laboratory for critical comments on this work and the National Institute of Health (R01GM107311-8) grant awarded to KD. Plasmid CMV-B- GECO1 is a gift from Dr. Robert Campbell (Addgene plasmid #32448); Plasmids pGP-CMV-GCaMP6f and pGP-CMV-NES-jRGECO1a are gifts from Dr. Douglas Kim (Addgene plasmid #40755 and #61563); Plasmids pCMV-G-CEPIA1er and pCMV-R- CEPIA1er are gifts from Dr. Masamitsu Iino (Addgene plasmid # 58215 and # 58216); Plasmid pME-pHluorin2 is a gift from Dr. David Raible (Addgene plasmid #73794).

